# Differential transcription of expanded gene families in central carbon metabolism of *Streptomyces coelicolor* A3(2)

**DOI:** 10.1101/849802

**Authors:** Jana K Schniete, Richard Reumerman, Leena Kerr, Nicholas P Tucker, Iain S Hunter, Paul R Herron, Paul A Hoskisson

**Affiliations:** Strathclyde Institute of Pharmacy and Biomedical Sciences, University of Strathclyde, 161 Cathedral Street, Glasgow, G4 0RE, UK; Isomerase Therapeutics, Cambridge, CB10 1XL, UK; Institute of Earth and Life Sciences, School of Energy, Geoscience, Infrastructure and Society, Heriot-Watt University, Riccarton, Edinburgh, EH14 4AS, UK

**Keywords:** *Streptomyces*, RNA-Seq, central carbon metabolism, metabolic plasticity, gene redundancy, silent biosynthetic clusters, metabolic engineering

## Abstract

**Background:** Streptomycete bacteria are prolific producers of specialised metabolites, many of which have clinically relevant bioactivity. A striking feature of their genomes is the expansion of gene families that encode the same enzymatic function. Genes that undergo expansion events, either by horizontal gene transfer or duplication, can have a range of fates: genes can be lost, or they can undergo neo-functionalisation or sub-functionalisation. To test whether expanded gene families in *Streptomyces* exhibit differential expression, an RNA-Seq approach was used to examine cultures of wild-type *Streptomyces coelicolor* grown with either glucose or tween as the sole carbon source.

**Results:** RNA-Seq analysis showed that two-thirds of genes within expanded gene families show transcriptional differences when strains were grown on tween compared to glucose. In addition, expression of specialised metabolite gene clusters (actinorhodin, isorenieratane, coelichelin and a cryptic NRPS) was also influenced by carbon source.

**Conclusions:** Expression of genes encoding the same enzymatic function had transcriptional differences when grown on different carbon sources. This transcriptional divergence enables partitioning to function under different physiological conditions. These approaches can inform metabolic engineering of industrial *Streptomyces* strains and may help develop cultivation conditions to activate the so-called silent biosynthetic gene clusters.

## Introduction

Streptomycete bacteria are major source of clinically useful bioactive natural products including antibiotics, immunosuppressive and anti-cancer agents. A remarkable feature of their genomes is that there are often several genes that appear to encode the same biochemical function (Bentley *et al*., 2002; Schniete *et al*., 2018). This is often referred to as ‘genetic redundancy’, where two or more genes are performing the same biochemical function (Nowak *et al*., 1997). While genetic redundancy does occur in nature, many so called ‘redundant’ genes have evolved divergent functions and provide the functional diversity and evolutionary robustness that can be observed in genomes from a wide range of organisms (Wagner, 2008b). There are two main mechanisms that contribute to the expansion of gene families within genomes - the duplication of genes or horizontal gene transfer events (Treangen & Rocha, 2011; Wagner, 2008a, b). Such gene family expansions are well known in streptomycete regulatory genes (Bush, 2018; Chater & Chandra, 2006; Clark & Hoskisson, 2011; Girard *et al*., 2013) and in specialised metabolite biosynthesis (Jenke-Kodama *et al*., 2005; Ridley *et al*., 2008). Yet, there has been limited attention focussed on the genes of central metabolism despite a number of gene expansion events being identified from biochemical and phylogenomic studies. There is an increasing appreciation that gene expansion events in central metabolism may facilitate the evolution of specialised metabolites (Borodina *et al*., 2008; Cruz-Morales *et al*., 2016; Fernández-Martínez & Hoskisson, 2019; Noda-García & Barona-Gómez, 2013; Noda-García *et al*., 2013; Schniete *et al*., 2018). Whilst expansion of gene families is widespread, expression differences, gene dosage, cofactor variation, allostery or substrate affinities are seldom taken into account during the construction of metabolic models for *Streptomyces*. As a result, homologous functions often combined into a single flux pathway that may not reflect the physiological nature of each gene product. This is especially stark when attempting to understand the supply of precursor molecules from central metabolism to the production of specialised metabolites.

It was hypothesised that gene families that have undergone gene expansion events would exhibit different expression profiles if they have diverged functionally. To understand the role of these gene expansions in *Streptomyces* and their impact on central metabolism and specialised metabolism, an RNA-Seq approach was taken using *S. coelicolor* grown either on glucose or tween as sole carbon source. Cultures were compared at a single point during growth (mid-log phase) to understand how transcription of central metabolic genes varies when the cultures are growing primarily via glycolysis (glucose as the sole carbon source) or gluconeogenically (with the mono-oleate, tween). It was found that expanded gene families exhibit different responses to growth on different carbon sources. These data will enable prioritisation of targets for metabolic engineering and will be informative for the construction of metabolic models.

## Results and discussion

### Expanded gene families in carbon metabolism exhibit different transcriptional profiles

To investigate the differences between glycolytic and gluconeogenic growth conditions, *S. coelicolor* M145 (an A3(2) strain) was grown respectively on glucose and Tween40 as sole carbon source. Growth of liquid cultures was monitored by cell dry weight and samples were removed for total RNA extraction at mid-exponential phase, which for glucose was 19 h and for tween was 36 h (Supp. Fig. 1). The specific growth rate of *S. coelicolor* M145 was 0.18 h^-1^ when grown on glucose and 0.4 h^-1^ for growth on tween (See Supp. Fig. 1).

RNA samples from these cultures showed global effects on transcription, when analysed by RNA-Seq, when the strain was grown on different carbon sources. Growth on tween resulted in 644 genes being differentially expressed when compared to growth on glucose. Of the 644 differentially expressed genes, 37% were predicted to encode hypothetical proteins, 9% encoded regulatory genes, 12% encoded central carbon metabolic enzymes, 6% were transporters, 4% were associated with specialised metabolism, 2% encoded genes associated with stress metabolism and 1 % of genes were associated with DNA replication, nitrogen metabolism and genes associated with metal metabolism (See Supp. Table 1-6). Growth on Tween40 resulted in an up-regulation of genes associated with gluconeogenesis and fatty acid degradation when compared to cultures grown on glucose (Fig. 1C and Supp. Tab.1-6).

**Figure 1).**
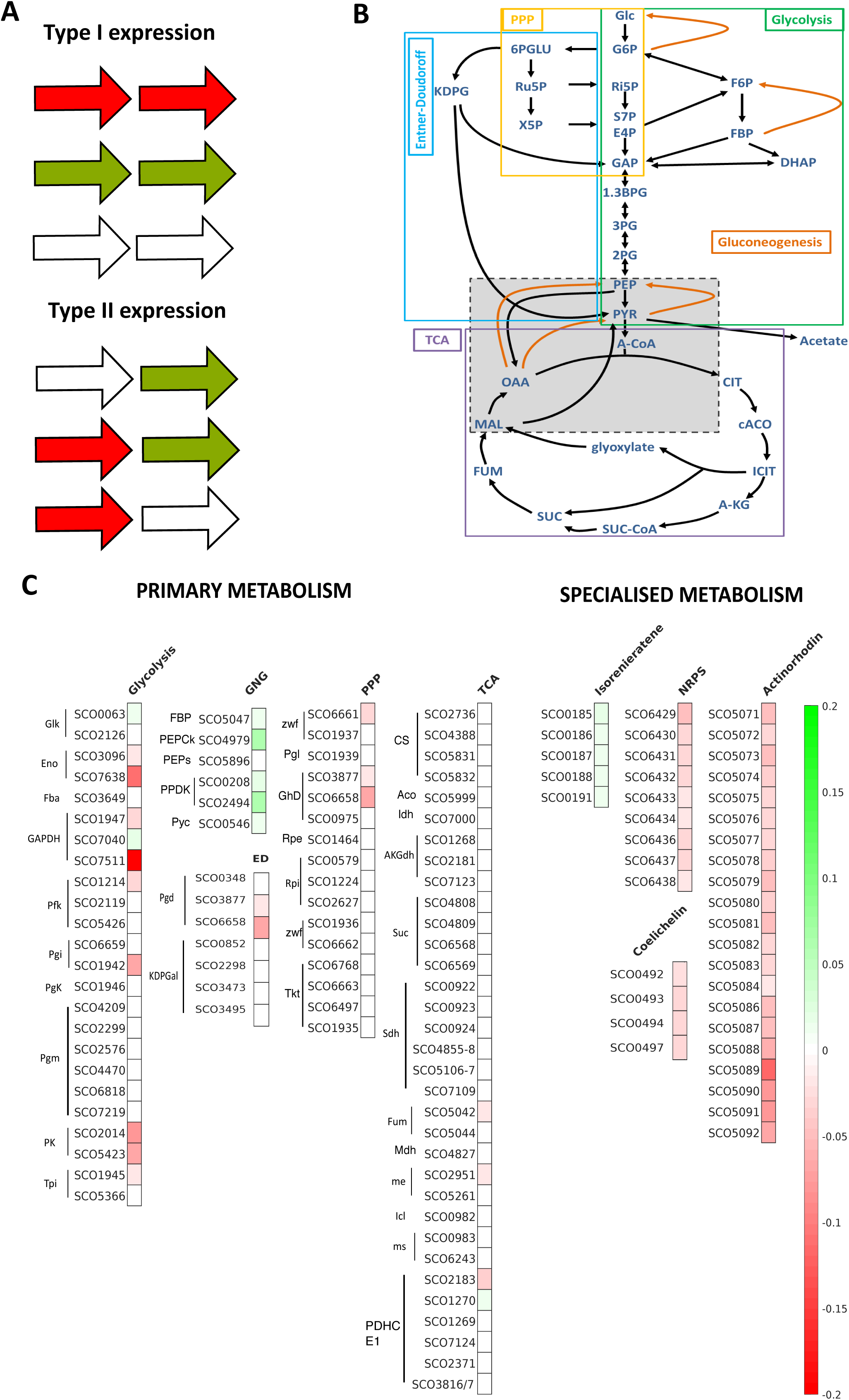
Carbon source-dependent expression of genes in *S. coelicolor*. **1A).** Schematic overview of expression patterns observed for expanded gene families Type I-expression where genes behaved in the same manner under the different growth conditions and Type II, where members of an expanded gene family exhibited differential gene expression across multiple gene family members (Green = up-regulated; Red = down-regulated & white = no expression change) **B)**. Schematic overview with main metabolites of the central carbon metabolism grouped according to pathways: glycolysis (green), pentose phosphate pathway (PPP, yellow), Entner-Doudoroff pathway (not complete in *S. coelicolor;* blue), tricarboxylic acid cycle (TCA, purple), gluconeogenesis (orange arrows). **C)**. Visualisation of gene expression using Heatmap by pathway (gene expression was normalised to maximum and minimum values of the entire dataset, and heat map colouring was based on maximum and minimum values of genes represented, where green is up-regulated and red is down-regulated)

To examine transcription of expanded gene families, two categories of expression profiles were considered: Type 1, where all genes within an expanded gene family behaved similarly under the different growth conditions, and Type II where members of an expanded gene family exhibit differential gene expression i.e. one or more member of the family increased, while expression of other members of the gene family, increased to a different degree, decreased or remained the same (Fig. 1A and Supp. Table 1). We identified 34 enzymatic reactions in central metabolism whose expression differed when cultures were grown on tween or glucose. Of these, 21 had more than one enzyme predicted to encode the same enzymatic function, i.e. were expanded gene families (Fig. 1; Supp. Table 1). When the expression profiles of these genes were examined further, 9/21 enzymatic reactions had families that showed Type I expression profiles and 12/21 expanded gene families exhibited Type II patterns of expression. This indicates that despite their identity in genome annotation, members of some expanded gene families are not redundant in function but have distinctive physiological roles (Borodina *et al*., 2008; Gubbens *et al*., 2012; Schniete *et al*., 2018). It is unlikely that functional differences can be attributed at the transcriptional level. However the data enable prioritisation of targets for metabolic engineering, such as modulating the expression poorly expressed members of gene families.

The glycolysis pathway, unsurprisingly, showed reduced expression when the cultures were grown on Tween40 as sole carbon source. Only two glycolytic enzyme families exhibited Type II expression, with an increase in expression for the minor glucose kinase (SCO0063; 5-fold increase in expression), whereas expression of the primary *glk* (SCO2126) remained unchanged under both conditions. GAPDH also showed Type II expression profiles, with expression of SCO7040 increased on tween, whereas the other two copies of the genes encoding GAPDH had reduced expression. A previous proteomic-based study also identified an increase in abundance of the SCO7040 protein when grown on a non-glucose carbon source (fructose; Gubbens *et al*., 2012). The two PK genes, as we have previously demonstrated, exhibited Type I expression (Schniete *et al*., 2018), with both copies being down regulated when cultures are grown on tween.

Overall, genes involved in gluconeogenesis were upregulated when cultures were grown on tween compared to glucose - as would be expected. The only expanded gene family encoding gluconeogenic function is that encoding the two PPDKs, which both displayed substantially increased expression (Type I expression), with *ppdk1* up regulated 11-fold up and *ppdk2* upregulated 30-fold (*ppdk2*). This suggests that, under these conditions, *S. coelicolor* may be using unconventional gluconeogenic routes for anaplerotic reactions rather than via the glyoxylate shunt, as expression of ICL and both MS enzymes remain unchanged during growth on either carbon source.

Expression of the pentose phosphate pathway showed little differential expression under the two conditions studied. Type II expression profiles were observed for *zwf*, with expression of SCO6661 reduced when cells were grown on tween, whereas expression of SCO1937 was unchanged between the two conditions. This was consistent with the work of Gubbens *et al*. (2012), where SCO6661 was downregulated when cultures were grown on fructose rather than glucose. There was no evidence of changes in transcriptional activity for the putative Entner-Doudoroff (ED) pathway genes for KDPG aldolase (SCO2298, SCO3473 and SCO3495). However reduced expression of two of the three phosphogluconate dehydratase homologues (SCO3877 and SCO6658) was observed when cultures were grown on tween (Fig. 1C). Whilst it was reported previously that there is no active ED pathway in *S. coelicolor* (Gunnarsson *et al*., 2004) these data suggest that putative phosphogluconate dehydratase homologs do respond to changes in carbon source.

In general, expression of genes encoding enzymes of the tricarboxylic acid (TCA) cycle remained stable under both growth conditions as would be expected, given this core part of metabolism is required for biosynthesis under all physiological conditions. The exception was expanded gene families, which exhibited differential expression (Type II): expression of the fumarase (SCO5042) was reduced two-fold on tween whilst SCO5044 expression remained unchanged; one copy of the malic enzyme (SCO2951) showed a two-fold decrease in expression, compared to SCO5261, which remained unchanged under both conditions; the extensive expansion in *S. coelicolor* of genes encoding the PHDC_E1_ subunit exhibited a range of expression changes - expression profiles for SCO1269, SCO7124, SCO2371 and SCO3816/3817 were the same for both carbon sources, whereas SCO2183 had a 3.5-fold decrease and SCO1270 a six-fold increase on tween.

As expected the fatty acid utilisation genes were expressed more when *S. coelicolor* was grown on tween, especially cholesterol esterase (SCO5420), which had an increase of almost 37-fold. The enzyme catalyses the hydrolysis of the head group of tween from the fatty acid palmitate, which are then utilised as carbon source (Plou *et al*., 1998; Pratt *et al*., 2000; Sakai *et al*., 2002). The other genes from this pathway showed between 3 to12-fold increase in expression (Supp. Table 2).

To verify the RNA-Seq data, we performed qPCR on the RNA samples used for the RNA-Seq experiment using primer pairs for five different genes - *pyk1* and *pyk2* from glycolysis, *ppdk1* and *ppdk2* from gluconeogenesis with primary sigma factor *hrdB* as control. These two pairs of genes were chosen as representative of expanded families, identified in a previous study (Schniete *et al*., 2018). RNA-Seq data showed that expression was strongly up or down-regulated under the chosen conditions, whilst *hrdB* (as expected) showed no difference under the two conditions tested. The fold-change difference in the qPCR data was similar to that observed in RNA-Seq experiments, corroborating the wider results (Supp. Fig. 2 & 3).

### Carbon source influences specialised metabolite gene expression

Genes involved in isorenieratene biosynthesis (SCO0185-0191), showed an increase in transcription in cultures grown on tween, with a three to eight-fold increase in expression across the entire operon. Isorenieratene is associated with blue light exposure, with the pathway present in green photosynthetic bacteria and a few actinobacteria (Krügel *et al*., 1999; Takano *et al*., 2005). Isorenieratene is a carotenoid with antioxidative properties; it is synthesized via the mevalonate-independent pathway (MEP/DOXP pathway) from basic precursors of GAP and pyruvate, derived from glycolysis to form Isopentenyl pyrophosphate (IPP) and Dimethylallyl pyrophosphate (DMAP). These metabolites then enter the carotenoid biosynthetic pathway via phytoene, lycopene and beta-carotene. The MEP/DOXP pathway also had increased expression of some genes (SCO6768 *-* 2.2-fold, SCO5250*-*4.4-fold) when grown on glucose (Supp. Table 2). Given the only difference between the cultures was the carbon source, we hypothesise that cultures experienced oxidative stress. This is supported by a seven-fold increase in expression of a superoxide mutase (SCO0999; Supp. Table 2).

A non-ribosomal peptide synthetase (NRPS) pathway (*SCO6429-6438*), the siderophore coelichelin biosynthetic gene cluster (SCO0491-0498) and the actinorhodin cluster (*SCO5071-5092*) all had decreased expression when grown on tween compared to glucose by two to five-fold, two to three-fold and three to 11-fold respectively (Fig. 1 and Supp. Table 2). This may reflect a tighter control of entry into specialised metabolism when cultures are grown on tween rather than glucose, although further experiments are needed to confirm this.

Studies such as this can inform metabolic modeling approaches for strain improvement enabling expression data and metabolic flux analysis to be taken into account with respect to isoenzymes within expanded gene families rather than combining the activities of all copies of a gene family into a single flux (Fernández-Martínez & Hoskisson, 2019). This knowledge will inform choices of metabolic engineering targets and the supporting dataset will also inform on the role of hypothetical proteins and regulators during growth on two different carbon sources that drive metabolism along two different pathways.

## Materials and Methods

### Bacterial strains and growth conditions

*Streptomyces coelicolor* A3(2) M145 (Kieser *et al*., 2000) was used throughout the study. Spores were germinated in 50 ml of 2x YT medium in flasks containing a metal spring and were incubated for up to 8h at 30°C and 250 rpm until emerging germ tubes were visible under a microscope (Kieser *et al*., 2000). The cultures were harvested by centrifugation, washed twice with 0.25 M TES buffer (pH 7.2) and pellets were resuspended in media. Growth curves were performed at 400 ml scale with minimal medium (Hobbs *et al*., 1989) in 2 L flasks containing a metal spring at 30°C, shaken at 250 rpm. Carbon source added was adjusted to ensure that each culture contained 166.5 mM equivalent of carbon.

### RNA extraction and sample preparation

*S. coelicolor* biomass (15 ml sample) from liquid cultures was harvested by centrifugation (5 min, 4°C, 4000 x *g*). Cell pellets were resuspended in an equal volume of RNAprotect (Qiagen) for 5 min at room temperature. Following centrifugation (5 min, 4°C, 6000 × *g*) biomass was then resuspended in 1ml 1x TE buffer containing 15 mg/ml lysozyme. Tubes were vortexed for 10 s and incubated at room temperature for 60 min whilst shaking. 1 ml RLT buffer (Qiagen RNA Isolation Kit) + 10 μl β-mercaptoethanol was added and vortexed and a phenol chloroform extraction followed by an ethanol precipitation was carried out. The sample was then purified using a commercial RNA isolation Kit (Qiagen). The isolated RNA was treated with RNAse free DNase (Ambion, Life Technologies) as specified by the manufacturer.

Quantification of RNA was carried out using Qubit® (Life Technologies). The quality and integrity of the RNA was assessed using a Bioanalyzer (Agilent). Furthermore, the RNA samples were also used as templates for a generic PCR in order to check for DNA contamination using primers for *hrdB* (SCO5820).

To enrich the samples for mRNA, rRNA depletion was performed (rRNA depletion Kit Ribo Zero Magnetic Kit for Gram-positive bacteria; Epicentre [Illumina]) according to the manufacturer’s instructions. The rRNA-depleted samples were then precipitated with ethanol and resuspended according to the manufacturer’s instructions. The quality and integrity of the samples were then analysed on the BioAnalizer (Agilent) and the concentration was determined using Qubit (Life Technologies).

### Library preparation

cDNA synthesis and library preparation was carried out using the Ion Total RNA-Seq Kit v2 Revision E from Ion Torrent, Life Technologies. The manufacturer’s protocol for less than 100 ng rRNA depleted samples was followed. The yield and size distribution of the amplified cDNA was assessed using BioAnalizer (Agilent). The three samples harvested from glucose cultures and three samples from tween grown cultures were barcoded and pooled to a concentration of 20 pM as specified by the manufacturer. The Ion OneTouch 2 system using the Ion PGM Template OT200 Kit (Life Technologies) was used for template preparation which includes the steps of emulsification, amplification and enrichment of the library. The libraries were checked using the quality control assay for the Qubit (Life Technologies).

### RNA Sequencing

Sequencing of the samples was carried out using a Ion Torrent Personal Genome Machine System (PGM; Life Technologies) on a 316v2 chip following the procedures in the manual, Ion PGM Sequencing 200 Kit v2 User Guide Revision 3.0. All sequence data were deposited on the Sequence Read Archive (SRA) under the BioProject: PRJNA566372 (https://www.ncbi.nlm.nih.gov/sra/PRJNA566372).

### Data analysis

Sequencing data were downloaded from the Ion Torrent server version Torrent Suite 4.0.2 in Fastq format. Reads were trimmed and low quality data removed. Reads were mapped to the reference genome of *S. coelicolor* (Bentley *et al*., 2002); Genbank: NC_003888.3). The data were analysed using CLC Genomics Workbench (Version 7.5, Qiagen). The software showed on average 99.8% and 99.7% alignment of the reads to the reference sequence. The CLC differential gene expression tool was used to determine differential gene expression. To examine differential gene expression analysis within the dataset, the cut-off was set at a p-value of 0.05. Genewise dispersions are estimated by conditional maximum likelihood using the total count for the gene of interest followed by empirical Bayes to obtain a consensus value (Smyth and Verbyla, 1996; Robinson and Smyth, 2007). The differential expression is then assessed using Fisher’s exact test adjusted to over-dispersed data (Robinson and Smyth, 2008). The raw data output from CLC Genomics Suite can be found in Supp Table 7, and a list of all differentially expressed genes is shown in Supp. Table 5. For the heatmap representation in Fig 1C, all differentially expressed genes were normalised to the maximum (+1) and minimum (−1) of all significantly different expressed genes and this can be found in Supp. Table 6. The code to create Fig 1C can be found in the Supp. Code 1 (https://doi.org/10.6084/m9.figshare.10008914.v2). The colour code was expressed relative to the highest and lowest expression change in the data shown ranging from green to red respectively.

### qPCR

qPCR was performed on the same samples as the RNA-Seq in order to provide independent confirmation of the data obtained. Each primer pair was tested using genomic DNA as template. cDNA was synthesized from the RNA samples using qPCRBIO cDNA synthesis Kit (PCR Biosystems) following the manufacturer’s instructions. Quantification of the cDNA was carried out using the QuantiFluor ssDNA system (Promega) with the Qubit Fluorometer (Life Technologies).

All cDNA samples were diluted to a concentration of 10 ng/μl and each reaction contained 10 ng of cDNA. Samples were mixed with qPCRBio MasterMix and the respective primers (Supp. Table 8; Kit 2x qPCRBIO SyGreen Mix Lo-ROX from PCRBIOSYSTEMS). Using a Corbett Research 6000 (Qiagen) machine, PCR reactions were subjected to a three-stage thermocycling reaction (one 3 min step at 95 °C, followed by 40 cycles of 5 s at 95 °C and one 25 s step at 60 °C. Each reaction was carried out in duplicate and a no template control was included for each set of primers. To allow quantification, standard curves for each gene were prepared (in triplicate) using purified PCR product from a genomic DNA PCR. This template was diluted to create seven different standards ranging from 10^1^-10^7^ molecules/per reaction. (Supplementary Figure 3) and were used to calculate the concentrations of the unknown samples obtained in the RNA Sequencing.

## Author Contributions

Conceptualization: JKS, PAH, NPT, LK

Data curation: JKS & RR

Formal analysis: JKS & RR

Funding acquisition: PAH, PRH, ISH

Methodology: JKS, RR, PAH, PRH, NPT, LK

Project administration: PAH & PRH

Supervision: PAH, ISH & PRH

Writing – original draft: JKS and PAH

Writing – review and editing: JKS, RR, LK, NPT, ISH, PRH, PAH

## Acknowledgements

The authors would like to thank Jan Schniete for writing the MATLAB code for the heatmap representation in Figure 1C. PAH would like to acknowledge the support of NERC (grant: NE/M001415/1) and BBSRC/NPRONET (grant: NPRONET POC045). PAH and PRH acknowledge SULSA funding for PhD studentships to JKS and RR.

## Ethical statement

No ethical approval was required.

## Legend for abbreviations

Catalytic functions:

Glk: glucose kinase
Zwf: Glucose-6-phosphate 1-dehydrogenase
Pgl: 6-phosphogluconolactonase
GhD: 6-phosphogluconate dehydrogenase
Rpi: Ribose-5-phosphate isomerase
Rpe: Ribulose 5-Phosphate 3-Epimerase
Tal: transaldolase
Tkt: transketolase
Pgd: 6-phosphogluconate dehydratase
KDPGal: KDPG aldolase
Eno: enolase
Fba: fructose-1,6-bisphosphate aldolase
Gap: glyceraldehyde-3-phosphate dehydrogenase
Pfk: phosphofructokinase
Pgi: Phosphoglucose isomerase
Pgk: Phosphoglycerate kinase
Pgm: Phosphoglycerate mutase
Pyk: pyruvate kinase
Tpi: triosephosphate isomerase
FBPase: FBP bisphosphatase
PEPCk: PEP carboxykinase
PPS: PEP synthase
PPDK: pyruvate phosphate dikinase
Pyc: Pyruvate carboxylase
CS: citrate synthase
Aco: aconitase
Idh: isocitrate dehydrogenase
AKGdh: Alpha-ketoglutarate dehydrogenase
Suc: succinyl-CoA synthetase
Sdh: succinate dehydrogenase
Fum: Fumarase
Mdh: malate dehydrogenase
me: malic enzyme
Icl: isocitrate lyase
Ms: malate synthase
PDHC: pyruvate dehydrogenase complex

**Metabolites**

Glc: Glucose
G6P: Glucose-6-Phosphate
6-PGLU: 6-phosphogluconate
Ru5P: Ribulose-5-phosphate
X5P: Xylose-5-Phosphate
KDPG: 2-keto-3-deoxy-6-phosphogluconate
F6P: Fructose-6-Phosphate
FBP: Fructose 1.6-bisphosphate
DHAP: dihydroxyacetone phosphate
Ri5P: Ribose-5-Phosphate
S7P: Seduheptulose-7-Phosphate
E4P: Erythrose-4-Phosphate
GAP: glyceraldehyde-3-phosphate
1.3BGP: 1.3-bisphosphoglycerate
3PG: 3-phosphoglycerate
2PG: 2-phosphoglycerate
PEP-: Phosphoenolpyruvate
PYR: Pyruvate
ACoA: Acetyl-CoA
Cit: Citrate
cAco: cisAconitate
ICit: Isocitrate
A-KG-: α-Ketoglutarate
SucCoA: Succinyl-CoA
Suc: Succinate
Fum: Fumarate
Mal: Malate
OAA: Oxaloacetate

## Supplementary Figure Legends

All supplementary data is available here https://doi.org/10.6084/m9.figshare.10008914.v2

**Supplementary Figure 1** Growth profiles of liquid *Streptomyces coelicolor* M145 cultures grown in glucose or tween with sampling point indicator (green arrow) at mid log phase. Means of three biological replicates are shown as *ln* of cell dry weight (CDW) over time. Each experimental condition was carried out with three biological replicates. Each data point is the mean of three independent experiments and an error bar represents the standard deviation for that data.

**Supplementary Figure 2**. Verification of fold change data from RNA-Seq and qPCR obtained from three biological replicates with standard deviation of five genes, one constitutively expressed gene (*hrdB*), two from glycolysis (*pyk1, pyk2*) and two from gluconeogenesis (*ppdk1, ppdk2*). Experimental details are in the Materials and Methods. Pyk = pyruvate kinase; PPDK = pyruvate phosphate dikinase; hrdB = housekeeping sigma factor

**Supplementary Figure 3** Standard curves of each gene for the quantification of transcripts in qPCR experiments A purified PCR product from a PCR for each gene from genomic DNA served as template (Supp. Table 8). This template was diluted to derive seven different standards ranging from 10^1^-10^7^ molecules. The standard curves were prepared in triplicate and used to calculate the concentration in the unknown samples, which were then compared to the results obtained in the RNA Sequencing. Pyk = pyruvate kinase; PPDK = pyruvate phosphate dikinase; HrdB = housekeeping sigma factor.

## Supplementary Table Legends

**Supplementary Table 1** Differential gene expression (DE) and expression category for growth on tween versus glucose in central carbon metabolism showing all genes annotated for the function Legend: green = up, red = down, yellow highlighted genes = expanded gene in Streptomyces, ‘-’ symbol indicates no significant change in expression detected, expression type meanings: I) same direction of change or no change in all genes II) different direction of change in all genes.

**Supplementary Table 2** Specialised metabolite gene clusters with individual genes showing differential expression for growth on tween and glucose, showing SCO number, gene function, fold change and p-value.

**Supplementary Table 3** Genes involved in fatty acid metabolism and EM-CoA pathway showing differential expression for growth on tween and glucose, showing SCO number, gene function, fold change and p-value.

**Supplementary Table 4** Regulatory genes showing differential expression for growth on tween and glucose showing SCO number, gene function, fold change and p-value.

**Supplementary Table 5** List of all genes showing differential expression for growth on tween and glucose showing SCO number, gene function, fold change and p-value.

**Supplementary Table 6** List of all genes showing differential expression for growth on tween and glucose showing SCO number, normalised fold change (+1 highest increase in expression to -1 highest decrease in expression), fold change and p-value.

**Supplementary Table 7** Raw data output from analysis of RNA-Seq data from CLC Genomics Workbench 7.5.

**Supplementary Table 8** Primers utilised for qPCR specifying for which gene, direction, sequence, melting temperature, amplicon size.

